# Tactile Sensitivity to the Frequency Spectrum of Complex Vibrotactile Signals

**DOI:** 10.1101/2023.11.10.566309

**Authors:** Thanh-loan Sarah Le, Gilles Bailly, Eric Vezzoli, Malika auvray, David Gueorguiev

## Abstract

Haptic interactions with objects induce complex vibrotactile signals that are central to tactile perception. Despite the broad literature on vibrotactile perception, there is surprisingly little knowledge of the sensory processing of complex tactile signals made of multiple frequencies. To fill this gap, the study reported here investigated the impact of the constitutive pure tones of a complex vibrotactile signal on its perception. Participants completed a 3-AFC task, in which they were asked to identify an odd signal among two complex references. The odd signal was created by removing one pure tone from the reference, which varied in spectral composition, harmonicity, and inter-frequency intervals. Each reference signal was made of either two, three, or four pure tones. The results revealed that the removed pure tone’s value as well as the inter-frequency interval play a significant role on participants’ performance whereas changes in harmonicity, complexity have little impact. In addition, the results showed that the smaller the ratio between the removed frequency and the lowest one of the reference signals, the better the participants’ capacity to identify the one with a missing tone. Overall, the detection of a missing tone in a complex signal predominantly depends on the tone’s frequency value of the removed signal suggesting a tactile sensitivity to the frequency spectrum of complex vibrotactile signals.

**NEW & NOTEWORTHY:** This research reports an investigation of the respective roles of frequency range, harmonicity, and complexity on human perception of vibrotactile signals. The results revealed that only pure tones that are close to the lower end of the frequency spectrum are noticed when they are missing. This finding sheds new light on the mechanisms of frequency selectivity in touch.

## Introduction

When a finger moves to explore a surface, the ensuing sliding contact elicits vibrations on the skin, which exhibit complex waveforms (1). These vibrational cues mediate the richness of tactile perception (2, 3) by endowing perceptual attributes like roughness, friction or hardness (4, 5). Moreover, temporal codes related to the complex vibrotactile signals elicited during touch have been suggested to mediate the perception of natural textures (6, 7). Extensive research within the realm of tactile sensation has delved into the domain of vibrotactile perception but our comprehension remains mostly limited to simple signals made of a single frequency. Research on such simple signals have shown that humans are sensitive to both frequency and amplitude variation, commonly associated with the psychological percepts of pitch (8, 9) and intensity (10, 11), respectively. Humans can reliably discriminate differences of 9% in the frequency of sinusoidal vibrations irrespective of the amplitude (12) and most studies observed that perception of differences in frequency mostly follows Weber’s law (13–16).

A complex signal is made of several pure tones, each of them with its own amplitude and frequency. The sum of these signals creates a specific waveform that is shaped by the interferences of these individual pure tones (Fig. 1). Research revealed that the perception of complex signals is influenced by the number of frequencies composing the signal. Notably, differentiating between two complex signals has been shown to be more difficult than for simple signals (16, 17). The stimulated tactile channel also exerts an impact on the perception of complex signals (18). Indeed, participants are more sensitive to phase shifts in complex signals that are composed of low frequencies (10 and 30 Hz), which target the rapidly adapting type 1 receptors, compared to high frequencies (100 and 300 Hz), which target Pacinian receptors. These results suggest that the Pacinian channel is insensitive to phase variations, while non-Pacinian channels encode them. The disparate sensitivity of tactile channels might also explain why simple signals, characterized by sinusoidal waveforms are perceptually distinct from square and sawtooth waveforms at a given frequency because the latter are accompanied by multiple harmonics (19). Interestingly, it has been found that the capacity to discriminate between pure tones made of distinct frequencies decreases when two to-be-compared signals are complexified with the addition of a signal at 100 Hz (16). Moreover, two vibrotactile signals can elicit identical perception even if they diverge in terms of amplitude and constitutive frequencies (12, 17). For example, a pure tone is not discriminated from a diharmonic vibrotactile signal when the pure tone is higher in amplitude or when it is larger than the fundamental frequency of the diharmonic signal due to sensory masking (12). It has also been shown that the pitch of a two-tones vibration can be matched with a simple pure tone that is closer in frequency to the lower of the two tones (20). Similar research showed that the pitch of a frictional texture containing two spatial frequencies can be perceptually matched by a simple sinusoidal texture of intermediate frequency (21).

**Fig. 1.**
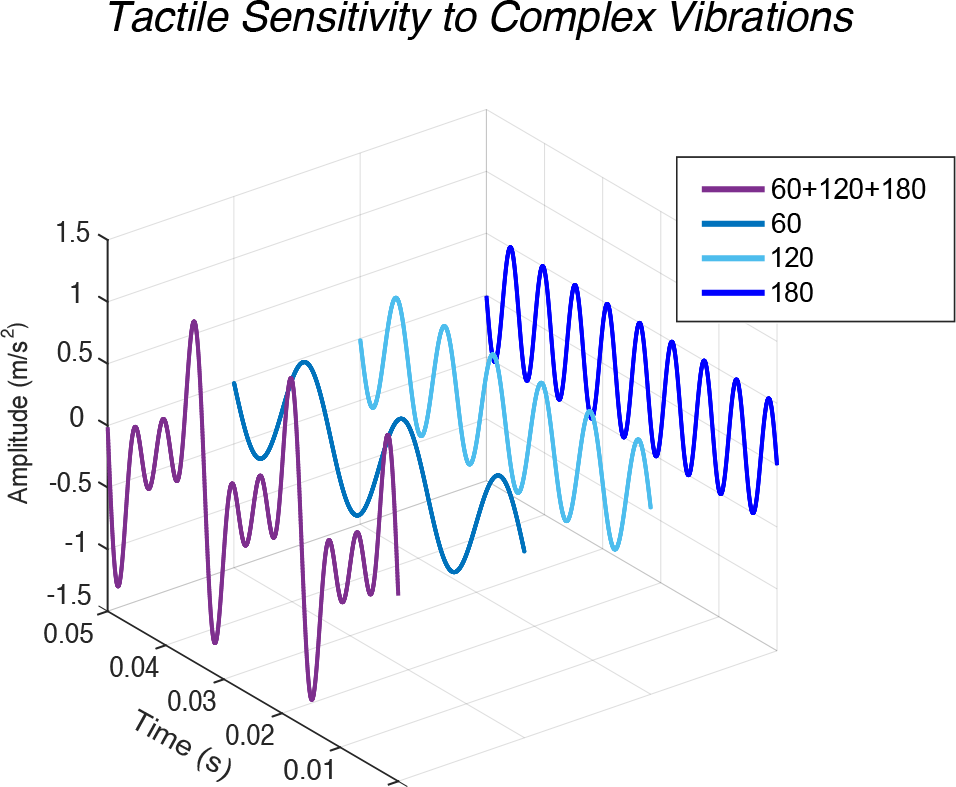
Example of a complex signal (60+120+180 Hz) obtained by superimposing sinusoidal waveforms at 60 Hz, 120 Hz and 180 Hz.

A complex signal is considered as harmonic when all its frequency components are integer multiples of its fundamental frequency and this characteristic has been investigated as a plausible cue for differentiating complex signals (19, 22, 23). Picinali et al. (22) found that the addition of a harmonic frequency to a simple sinusoidal signal is better discriminated than the addition of a non-harmonic frequency. This work suggested that non-harmonic sequences inherently yield greater discriminability, probably due to the emergence of amplitude beats. Russo et al. (23) showed that humans are capable to differentiate between cello, piano, and trombone tones played on their back with matching fundamental frequencies but instrument-specific harmonics. These studies highlight the influence of the vibrotactile signal’s harmonic content on its perception by humans.

The abovementioned studies have investigated the influence of frequency, amplitude, and harmonic content on tactile perception, however they mainly focused on signals composed of at most two frequencies. In addition, most of them only featured signals above 100 Hz, hence predominantly in the range of the Pacinian channel. Although a few models have been proposed (16, 17, 24) no general explanatory mechanism of the human perception of complex vibrations has emerged. In this context, the present study aims to investigate the detection of a missing pure tone from complex signals by implementing conditions that vary signals’ harmonicity, distances between frequencies within the signal (inter-frequency interval), and frequency ranges of the pure tones composing the signal. To that end, two psychophysical experiments were conducted with complex vibrotactile signals made of two, three, or four sinusoidal signals. The first experiment compared harmonic and non-harmonic signals. Frequencies in that experiment activate both the non-Pacinian and Pacinian channels. The second experiment investigated the influence of the inter-frequency interval within complex signals and implemented signals that predominantly activate the Pacinian channel. Thereby, a large number of diverse complex signals were tested in the two experiments, which enabled a subsequent analysis of the ratio between the removed pure tone and the lowest frequency of the signal as a potential sensory cue.

## Materials and Methods

### Participants

Two experiments were conducted separately in time (∼5 months). The procedure, materials, and methods were the same for both experiments but the vibrotactile signals were different. 12 participants (3 women and 9 men, mean age 25.3 yr, SD = 3.5) completed Experiment 1 and 12 participants (2 women and 10 men, mean age 26.6 yr, SD = 2.8) completed Experiment 2. All participants from the first experiment were contacted for participating in the second experiment and 8 of them agreed to do so. Participants were compensated 20 euros to complete each experiment, which took approximately 2 hours. The experiments were approved by the Research Ethics Committee at Sorbonne Université under approval CER-2021-104 and were conducted in accordance with the Declaration of Helsinki.

### Apparatus

An actuator, i.e., a voice-coil vibrator specifically tuned for haptic stimulation (Tactuator MM3C-HF, TactileLabs Inc.), delivered the various vibrotactile signals used in this study. The vibrotactile signals consisted in .wav files generated by a MATLAB (MathWorks, Inc.) script and they were verified by a Fast Fourier Transform (Fig. 2A).

**Fig. 2.**
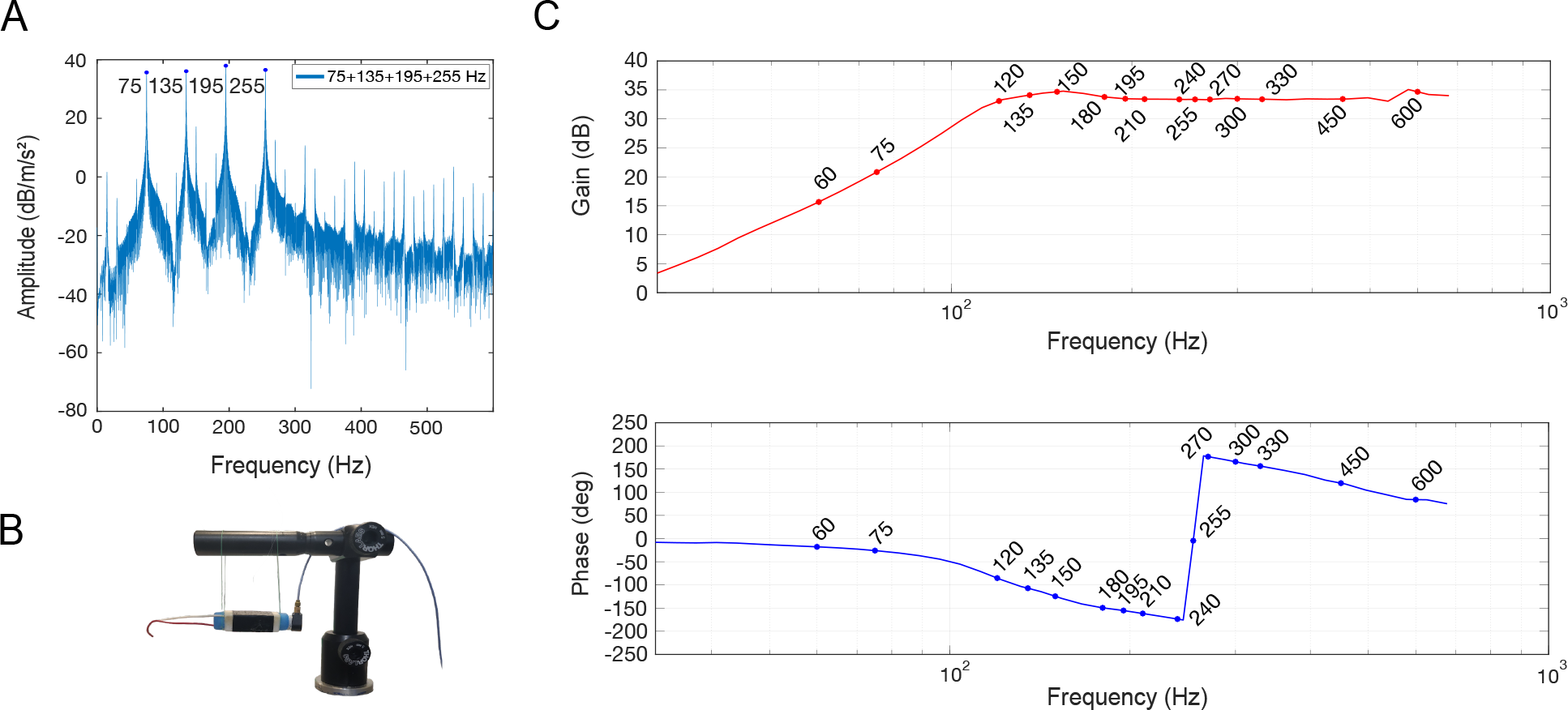
**A**) Fast Fourier Transform of the complex signal 75+135+195+255 Hz. **B**) Actuator with the accelerometer glued to one lateral side is suspended by nylon strings. **C**) Bode diagrams representing gain and phase of the vibrations delivered by Tactuator MM3C-HF as a function of the delivered frequency.

Vibrotactile pure tones were equalized to reach a peak acceleration amplitude around 7 m/s^2^, which is well above the human vibrotactile threshold (25) by estimating a Bode diagram expressing the acceleration amplitude at each frequency. The Bode diagram was obtained by measurements from an accelerometer (PCB Piezotronics Accelerometer 352A24) attached to the actuator and hung by nylon strings to avoid resonant coupling (Fig. 2B, C). The signal delivered by the actuator shifted in phase depending on the frequency due to the magnetic characteristics of the actuator (Fig. 2C). No distortions of the signal were observed after acceleration measurements on the actuator.

### Stimuli

In each experiment, participants were presented with either a *reference* vibrotactile signal made of 2, 3 or 4 tones or a *target* vibrotactile signal that is identical to the reference one except for the removal of one of its pure tones (see Table 1, upper table). Given the 44 reference signals for the two experiments, 56 different target signals can be derived, resulting in 112 different possible comparisons between a reference and a target: 4 tones vs. 3 tones (16 comparisons), 3 tones vs. 2 tones (48 comparisons), 2 tones vs. 1 tone (48 comparisons). For more clarity, these comparisons are called “Four-tones’’, “Three-tones’’, and “Two-tones’’ respectively in line with the number of pure tones in the reference. As an example, the reference three-tones complex signal 75+195+255 Hz can only be compared with three two-tones targets 75+195 Hz, 75+255 Hz, and 195+255 Hz (see Fig. 3A). Table 1 summarizes all the signals used in the study.

**TABLE 1.** Stimuli used for each experiment. The upper table displays comparisons and the lower table displays references and target signals.

| Comparisons |  |  |  |  |  |
| --- | --- | --- | --- | --- | --- |
| Two-tones |  | Three-tones |  | Four-tones |  |
| Reference | Target | Reference | Target | Reference | Target |
| 2-tones | 1-tone | 3-tones | 2-tones | 4-tones | 3-tones |

|  | Condition | Signals |  |  |  |
| --- | --- | --- | --- | --- | --- |
|  |  | 1-tone | 2-tones | 3-tones | 4-tones |
| Experiment 1:<br>Harmonicity | HARMONIC | 60<br>120<br>180<br>240 | 60+120<br>60+180<br>60+240<br>120+180<br>120+240<br>180+240 | 60+120+180<br>60+120+240<br>60+180+240<br>120+180+240 | 60+120+180+240 |
|  | NON-HARMONIC<sup>1</sup> | 75<br>135<br>195<br>255 | 75+135<br>75+195<br>75+255<br>135+195<br>135+255<br>195+255 | 75+135+195<br>75+135+255<br>75+195+255<br>135+195+255 | 75+135+195+255 |
| Experiment 2:<br>Inter-frequency<br>interval | SMALL-INTER<sup>2</sup> | 150<br>210<br>270<br>330 | 150+210<br>150+270<br>150+330<br>210+270<br>210+330<br>270+330 | 150+210+270<br>150+210+330<br>150+270+330<br>210+270+330 | 150+210+270+330 |
|  | LARGE-INTER<sup>2</sup> | 150<br>300<br>450<br>600 | 150+300<br>150+450<br>150+600<br>300+450<br>300+600<br>450+600 | 150+300+450<br>150+300+600<br>150+450+600<br>300+450+600 | 150+300+450+600 |

<sup>1</sup>In the NON-HARMONIC level, the set of pure tones is identical to HARMONIC except that 15Hz were added to each of pure tones to make the stimuli non-harmonic.
<sup>2</sup>The distance between two consecutive pure tones is either 60Hz or 150Hz.

**Fig. 3.**
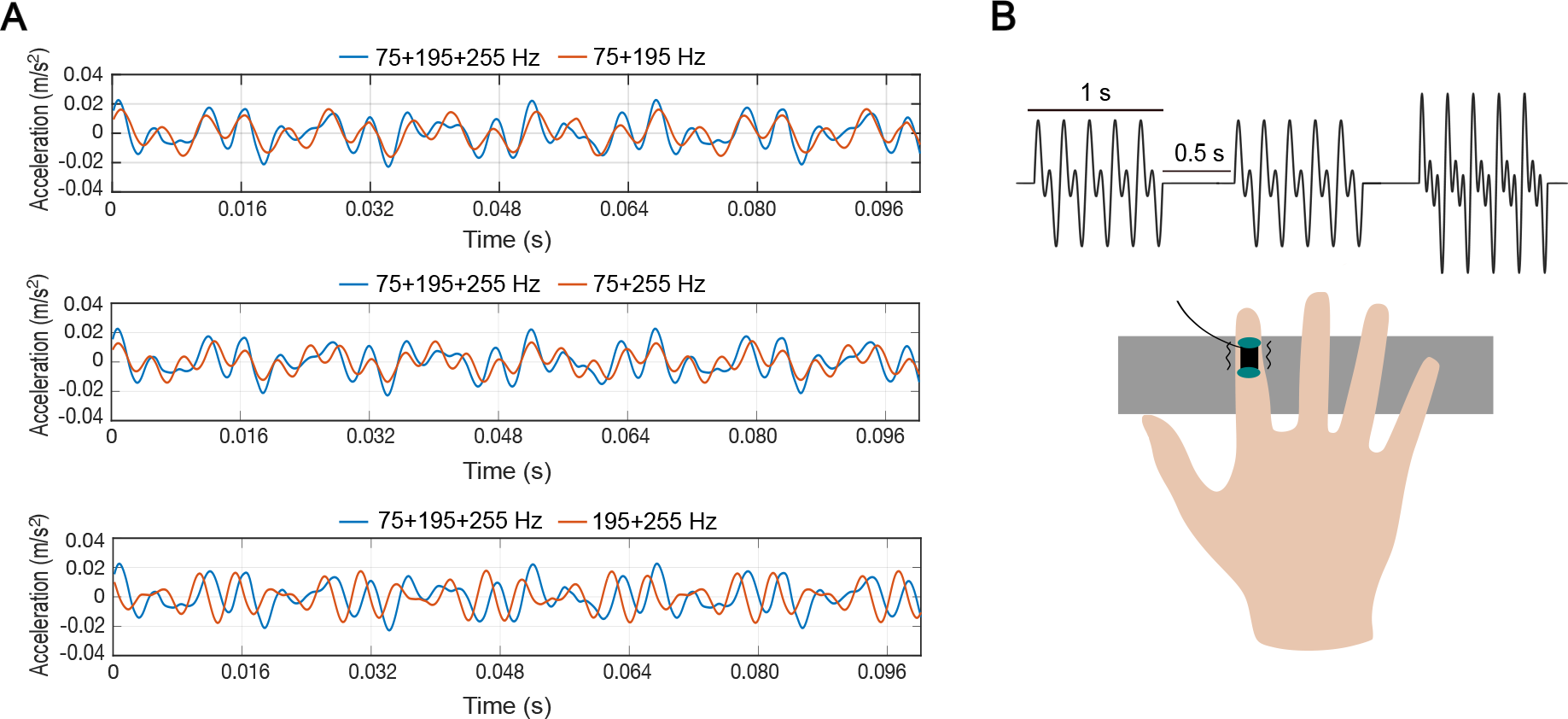
**A)** Example of the Three-tones vs. Two-tones comparisons with a vibrotactile reference comprising 75, 195 and 255 Hz. The blue curves represent the acceleration measure of the reference, and the red curves represent the possible target stimuli. **B)** The tactuator is placed over on the top of the intermediate phalanx of the right hand’s index finger and the participant performs a 3-AFC task that consists in reporting which signal out of three consecutives ones is different.

### Experimental procedure

In the two experiments, the participants performed a three-Alternative Forced Choice (3-AFC) task, which has the advantage to not force the experimenter to establish a dimensionality of interest prior to the experiment. The participants were comfortably seated with noise-canceling headphones that masked extraneous noise. The actuator was placed on the top of the intermediate phalange of their right hand’s index finger, attached with a plaster (Fig. 3B). The finger rested on foam to be more comfortable. Moreover, the actuator was not touching the nail to avoid undesired resonant frequencies. Once the participants read and signed the consent form, they completed six training trials to familiarize themselves with the task. In order to not influence the main experiment, the reference signals used during the training were built on different pure tones (50, 80, 130, 200 Hz). Data collection followed immediately after the training session. Each trial consisted of a sequence of three 1-second stimuli separated by a 0.5-second interstimulus interval. The three signals comprised two identical reference signals and an odd signal that was the target signal. The participants’ task was to identify which signal out of three was different from the other two (Fig. 3B). The participants were asked to indicate the position of the odd signal by pressing either 1, 2, or 3 on a keyboard followed by ‘Enter’ with no time limit. Once the answer was validated, a new trial started after 0.5 seconds. The participants could replay the stimulus only if they missed it. The presentation order of the three signals was randomized within each trial.

For each experiment, the presentation order of each condition level (Experiment 1: HARMONIC and NON-HARMONIC; Experiment 2: Small-inter and Large-inter) was counterbalanced across participants. Each condition included 28 unique comparisons, presented 8 times each. The 224 total trials per condition (28 comparisons x 8 repetitions) were presented in pseudo-randomized order. In addition to the 2 breaks given to the participant during a condition (every 74 trials), the participant was given a 15-minute break between the two conditions.

### Dependent and Independent variable

For experiments 1 and 2, the dependent variable is *Accuracy* corresponding to the correct answer rate (%) in the 3-AFC task. Both experiments also have in common two independent variables: *Reference* and *Removed pure tone. Reference*, from which the signal with the removed pure tone has to be discriminated, has 11 different levels whose values depend on the condition as illustrated in Table 1. *Removed Pure tone* indicates which pure tone is removed from the reference signal to create the target signal to be compared with. The different possible levels of *Removed Pure tone* depend on the number of constitutive frequencies of the reference signal. In addition, experiments 1 and 2 differed by the third independent variable that was considered (i.e., *Harmonicity* and *Inter-frequency interval*).

#### Experiment 1: harmonicity

The main independent variable is *Harmonicity* with two levels: HARMONIC and NON-HARMONIC. HARMONIC contains a set of pure tones (60, 120, 180 and 240 Hz) in harmonic sequence. In the NON-HARMONIC level, 15Hz are added to each pure tone in the harmonic set to make the stimuli non-harmonic.

#### Experiment 2: inter-frequency interval

The main independent variable is *Inter-frequency interval*, which is the distance between the four pure tones on which the reference signals were built. Two sets of pure tones were used in this experiment: 150, 210, 270, 330 Hz that are spaced by 60 Hz as in experiment 1 (Small-inter) and 150, 300, 450, 600 Hz that are spaced by 150 Hz (Large-inter). Since the created references result from all possible combinations of the four constitutive frequencies, inter-frequency intervals vary within a condition. Nonetheless, intervals in the Large-inter condition are generally larger than those in the Small-inter condition.

### Data analysis

The data were analyzed using RStudio software (version 4.1.3). The impact of removed pure tones from reference complex signals was analyzed by implementing a Generalized Linear Mixed Model (GLMM) on individual data. Due to the differences in the number of possible pure tones combinations, the Two-tones, Three-tones, and Four-tones comparisons were analyzed separately. D’Agostino and Pearson tests showed that the data were not normally distributed. Consequently, GLMM were fit by the maximum likelihood (Laplace Approximation) with logit link and binomial error distribution. Akaike Information Criterion Corrected weight (AICcWt), which is the proportion of the total amount of predictive power provided by the full set of models including independent variables, was assessed to select the better GLMM. Either the regression model included interaction terms between variables (i.e., one independent variable is influenced by another independent variable), or the model was simple (i.e., without interaction, each independent variable acts on the dependent variable). Independent variables were tested by analysis of deviance Type II Wald Chi-square tests. Post-hoc analyses were conducted on significant fixed effects between estimated marginal means by using the package ‘emmeans’ of Rstudio and responses were back-transformed from the logit scale (probability scale). Pairwise comparisons were adjusted by Holm’s method. In the additional analyses on the pure tones’ frequency, the method of least squares was used to fit data. AICc, RMSE and R-squared as well as F-test were computed to choose the best fit (High R-squared and Low AICc was preferred). AICc was chosen instead of AIC due to a small number of observations (26). According to Akaike’s findings (27) an AIC difference ≥2 should be a reference to select the best model, which also applies to AICc.

## Results

### Experiment 1: harmonicity

#### Two-tones

A Wald test conducted on the GLMM (*Reference, Removed pure tone, and Harmonicity*) showed a significant main effect of *Reference* on *Accuracy* (χ2(5) = 79.8, *P* < 0.0001). In contrast, the analysis yielded no significant main effect of *Harmonicity* on *Accuracy* (χ2(1) = 1.239, *P* = 0.266). Since the results for corresponding harmonic and non-harmonic references were very similar, they were merged for the subsequent post-hoc analyses. A posthoc Holm’s test on *Reference* revealed that pure tones removal from the references 180+240/195+255 Hz (M = 85.2%, SD = 35.6) was significantly more accurately detected than for any other reference (n = 2304, *P*-values < 0.01). On the other hand, pure tone removal from the 120+240/135+255 Hz (M = 59.4%, SD = 49.2) references was significantly less detected than for any other reference (n = 2304, *P*-values < 0.05). In addition, the Wald test showed a significant main effect of *Removed pure tone* on *Accuracy* (χ2(1) = 107.1, *P* < 0.0001). Post-hoc statistical tests on *Removed pure tone* showed that the removal of the lowest pure tone (M = 87.8%, SD = 32) was overall significantly more accurately detected than when the highest pure tone (M = 55.9%, SD = 49) was removed (Holm’s test, n = 2304, *P* < 0.0001). Finally, a significant interaction was found between *Removed pure tone* and *Reference* (χ2(5) = 118.5, *P* < 0.0001). Indeed, post-hoc tests on this interaction showed that the difference between the removal of the lowest and highest pure tone is not significant for the 120+180/135+195 Hz references (Holm’s test, n = 2304, *P* < 0.05) unlike for the other comparisons (Fig. 4A). This indicates that the reference complex signal from which a pure tone is removed impacts how sensitive humans are to its removal.

**Fig. 4.**
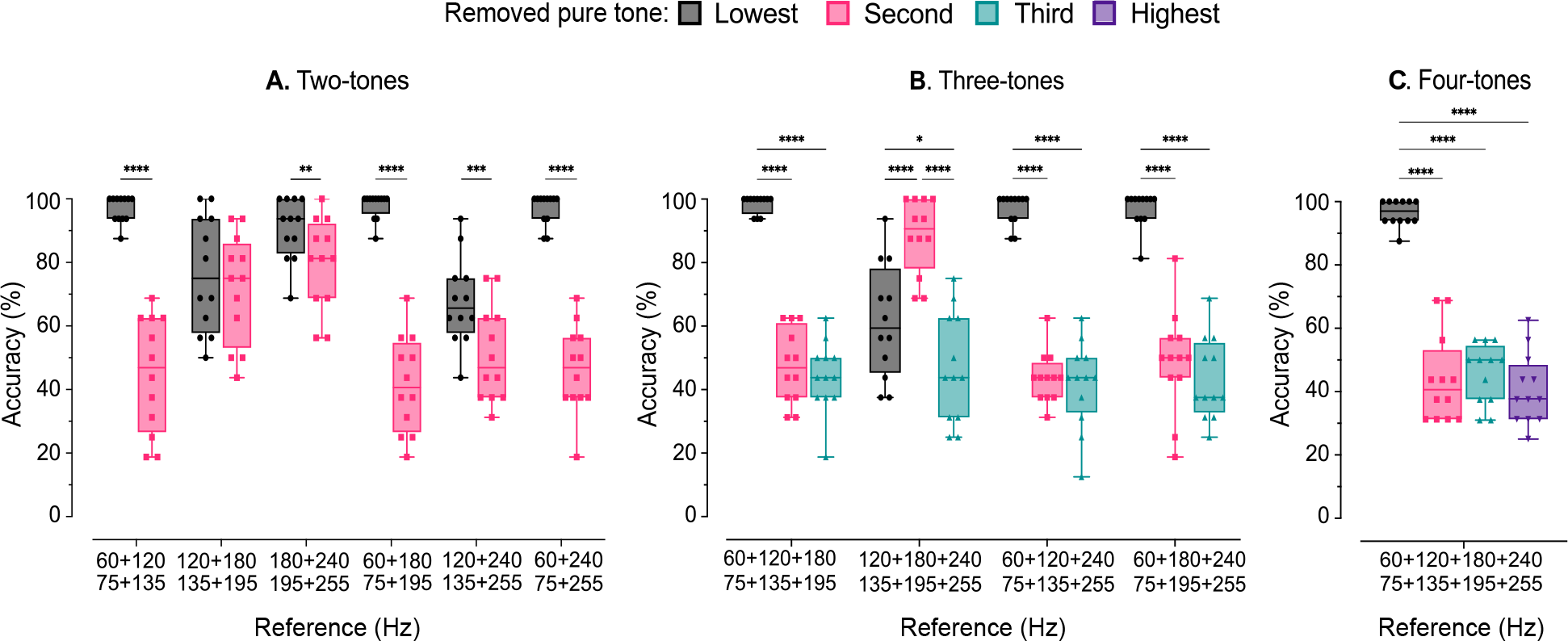
Accuracy (%) (median, interquartile range, and minimum to maximum) averaged across participants’ individual data for each combination of removed pure tone and reference signal. The HARMONIC and NON-HARMONIC conditions were merged since they showed no difference. **A)** References made of two pure tones. **B)** References made of three pure tones. **C)** References made of four pure tones. Asterisks represent statistical differences: * *P* < 0.05, ** *P* < 0.01, *** *P* < 0.001, **** *P* < 0.0001.

#### Three-tones

A Wald test conducted on the GLMM (*Reference, Removed pure tone*, and *Harmonicity*) showed a significant main effect of *Removed pure tone* on *Accuracy* (χ2(2) = 77.5, *P* < 0.0001). Moreover, a post-hoc Holm’s test on *Removed pure tone* showed that target signals with the lowest pure tone removed (M = 88.4%, SD = 32) were significantly more accurately discriminated from the reference than those with the second pure tone removed (M = 57.2%, SD = 49.5), which were in turn significantly more accurate than those with the highest pure tone removed (M = 44%, SD = 49.7) (n = 2304, *P*-values < 0.0001). Neither *Harmonicity* (χ2(1) = 0.6, *P* = 0.429) nor *Reference* (χ2(3) = 1.9, *P* = 0.584) had a main effect on *Accuracy*. A significant interaction effect was observed between *Removed pure tone* and *Reference* (χ2(6) = 189.7, *P* < 0.0001) indicating an effect of reference on how the removal of pure tones is perceived (Fig. 4B). Interestingly, removing the second pure tone was significantly better detected for the 120+180+240/135+195+255 Hz references than the removal of the highest pure tone (Holm’s test: n = 2304, *P* < 0.0001).

#### Four-tones

Only a Wald test with *Removed pure tone* and *Harmonicity* as independent variables was performed since four-tones signals feature only one reference. The Wald test conducted on the GLMM (*Removed pure tone*, and *Harmonicity*) showed a significant main effect of *Removed pure tone* on *Accuracy* (χ2(3) = 82.9, *P* < 0.0001). However, no significant main effect of Harmonicity on Accuracy was found (χ2(1) = 3.02, *P* = 0.082). Post-hoc pairwise comparisons performed on *Removed pure tone* revealed that participants discriminated significantly better a signal from its reference when the lowest pure tone was removed compared to the three other possible pure tones (Holm’s test: n = 768, *P*-values < 0.0001) (Fig. 4C).

Overall, the obtained results suggest that, in most cases, participants were more accurate at discriminating modified signals from their reference when the lowest pure tone is removed rather than when other pure tones are removed. Interestingly, this phenomenon did not occur for two reference signals starting with pure tones of 120/135 Hz. Moreover, no influence of the harmonic nature of the signal on the human capacity to detect the removal of a pure tone was observed. All effects of Experiment 1 are depicted in Table 2.

**TABLE 2.** Wald chi-square tests in Experiment 1.

| | Independent variables | $\chi^2$ | p-value |
| --- | --- | --- | --- |
| **Two-tones**<br>REFERENCE × REMOVED PURE TONE<br>+ HARMONICITY | REFERENCE | 79.8 | $P < 0.0001$ |
| | REMOVED PURE TONE | 107.1 | $P < 0.0001$ |
| | REMOVED PURE TONE × REFERENCE | 118.5 | $P = 0.0001$ |
| **Three-tones**<br>REFERENCE × REMOVED PURE TONE<br>+ HARMONICITY | REMOVED PURE TONE | 77.5 | $P < 0.0001$ |
| | REMOVED PURE TONE × REFERENCE | 189.7 | $P < 0.0001$ |
| **Four-tones**<br>REMOVED PURE TONE + HARMONICITY | REMOVED PURE TONE | 82.9 | $P < 0.0001$ |

The first column corresponds to the Generalized Linear Mixed Model including simple (+) or interaction ( × ) terms. Chi-square ( $\chi^2$ ) and $P$ -value are obtained by Wald Chi-square tests performed on independent and dependent variables (Accuracy).

### Experiment 2: Inter-frequency interval

In Experiment 2, the change of the inter-frequency interval also had a large impact on the frequency composition of the reference signals. Thus, independence of the reference and inter-frequency variables could not be assumed. Therefore, a first GLMM was performed with *Removed pure tone*, and *Inter-frequency interval* as independent variables. Subsequent GLMM statistical analyses, in which independent variables were *Removed Pure Tone* and *Reference*, were performed separately for the Small-inter and Large-inter conditions.

#### Two-tones

A Wald test conducted on the GLMM (*Removed pure tone*, and *Inter-frequency interval*) showed a significant main effect of *Removed pure tone* on *Accuracy* (χ2(1) = 310.9, *P* < 0.0001). A post-hoc Holm’s test on *Removed pure tone* revealed that, overall, participants better discriminate a signal from its reference when the lowest pure tone (M = 90.5%, SD = 29.4) was removed compared to the highest pure tone (M = 55.8%, SD = 49.7) (n = 2304, *P* < 0.0001). The Wald test also showed a significant main effect of *Inter-frequency interval* on *Accuracy* (χ2(1) = 48.8, *P* < 0.0001). A post-hoc Holm’s test on *Inter-frequency interval* revealed that participants better discriminate signals in the Small-inter condition (M = 79.0%, SD = 40.8) compared to the Large-inter condition (M = 67.3%, SD = 46.9) (n = 2304, *P* < 0.0001). Differences between Small-inter and Large-inter conditions as a function of *Removed pure tone* are depicted in Fig. 5A.

**Fig. 5.**
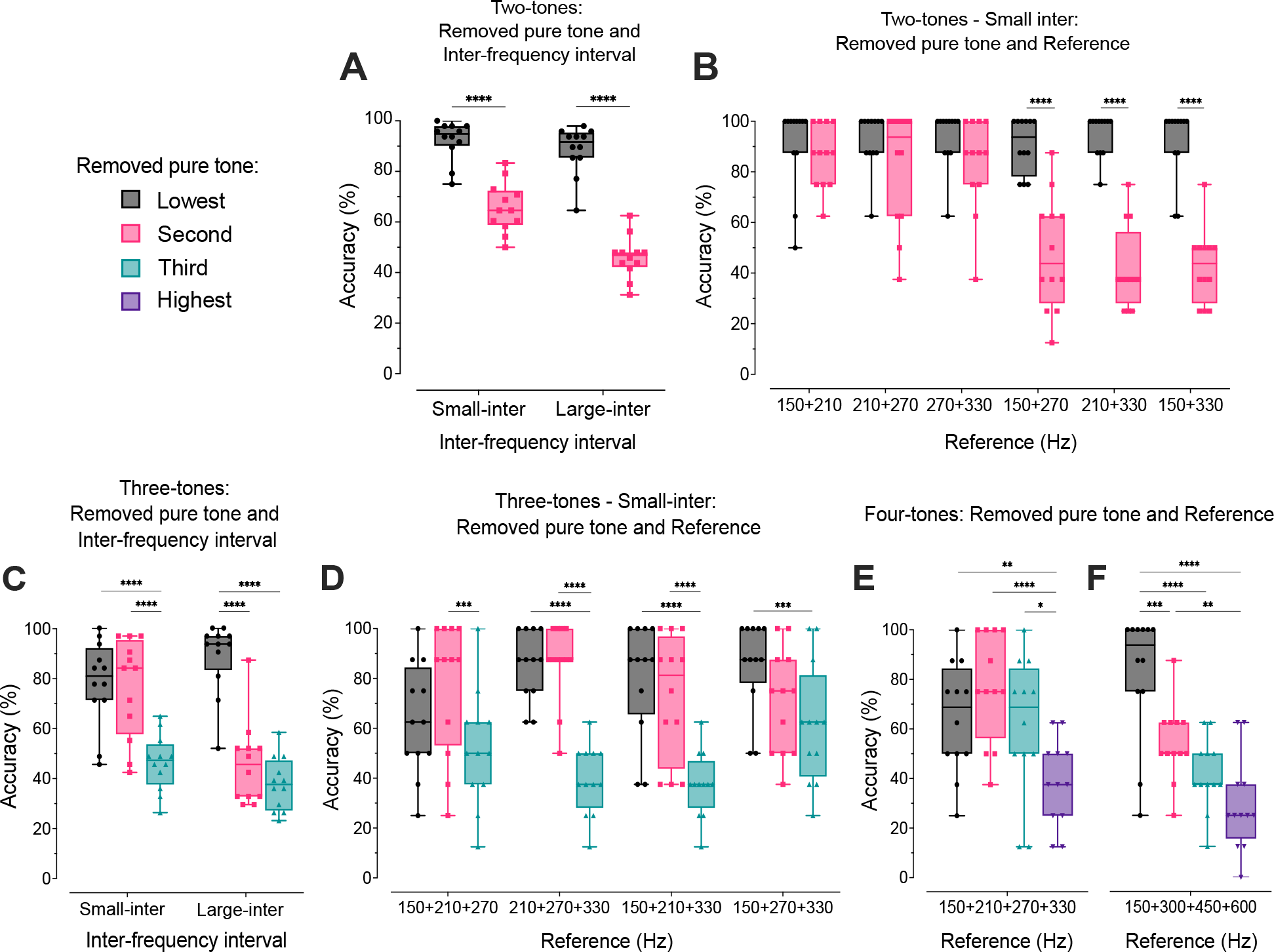
Accuracy (%) (median, interquartile range, and minimum to maximum) averaged across participants’ individual data as a function of **A**) Removed pure tone and Inter-frequency interval for the Two-tones comparisons, **B)** Removed pure tone and Reference in Small-inter for the Two-tones comparisons, **C)** Removed pure tone and Inter-frequency interval for the Three-tones comparisons, **D)** Removed pure tone and Reference in Small-inter for the Three-tones comparisons, **E)** Removed pure tone in Small-inter for the Four-tones comparisons, **F)** Removed pure tone in Large-inter for the Four-tones comparisons. Asterisks represent statistical significance: * *P* < 0.05, ** *P* < 0.01, *** *P* < 0.001, **** *P* < 0.0001.

These effects were further analyzed with a Wald test performed on both the Small-inter and Large-inter conditions (GLMMs: *Removed pure tone*, and *Reference*). In the Small-inter condition (Fig. 5B), the Wald test confirmed the significant main effect of *Removed pure tone* (χ2(1) = 92.1, *P* < 0.0001) and showed an effect of *Reference* (χ2(5) = 58.5, *P* < 0.0001) on *Accuracy*. A significant interaction effect was observed between *Removed pure tone* and *Reference* (χ2(5) = 27.1, *P* < 0.0001). A post-hoc Holm’s test on this interaction revealed that when the highest pure tone is removed, the participants’ discrimination accuracy is significantly higher for signals when the distance between the lowest and highest pure tone was smaller than 60 (Global mean 84%, SD = 36.7) than when this distance was larger than 60 (Global mean 47.2%, SD = 50) (n = 1152, *P*-values < 0.0001) (Fig. 5B). In the Large-inter condition, the Wald Test confirmed the significant main effect of *Removed pure tone* on *Accuracy* (χ2(1) = 188.6, *P* < 0.0001), as well as a significant interaction effect between *Removed pure tone* and *Reference* (χ2(5) = 12.6, *P* = 0.028). However, no significant effect of *Reference* was observed (χ2(5) = 1.39, *P* = 0.92). A post-hoc Holm’s test on *Removed pure tone* revealed that participants better discriminate a signal from its reference when the lowest pure tone was removed compared to the highest pure tone (n = 1152, *P* < 0.0001), as shown on Figure 5A.

#### Three-tones

A Wald test conducted on the GLMM (*Removed pure tone*, and *Inter-frequency interval)* showed a significant main effect of *Removed pure tone* on *Accuracy* (χ2(2) = 218.1, *P* < 0.0001). A post-hoc Holm’s test on *Removed pure tone* revealed that target signals with the lowest pure tone removed (M = 83.9%, SD = 36.8) were significantly easier to discriminate than signals with the second pure tone removed (M = 62%, SD = 48.6), which were significantly easier to discriminate than those with the highest pure tone removed (M = 43.5%, SD = 49.6) (n = 2304, *P*-values < 0.0001). The Wald test also showed a significant main effect of *Inter-frequency interval* on *Accuracy* (χ2(1) = 22.8, *P* < 0.0001). A post-hoc Holm’s test on *Inter-frequency interval* revealed that participants were more accurate in the Small-inter condition (M = 67.7%, SD = 46.8) than in the Large-inter condition (M = 58.5%, SD = 49.3) (n = 2304, *P* < 0.0001). In addition, the Wald test on the GLMM showed a significant interaction effect between *Removed pure tone* and *Inter-frequency interval* (χ2(2) = 69.9, *P* < 0.0001). Post-hoc Holm’s tests on this interaction indicated that the effect of the removal of the second pure tone depends on the interval type (Fig. 5C). Indeed, in Small-inter condition, signals with the second pure tone removed were significantly better discriminated than signals with the highest pure removed (n = 2304, *P*-values < 0.0001). In the Large-inter condition, these signals were significantly less accurately discriminated than the signals with lowest pure tone removed (n = 2304, *P*values < 0.0001). Overall, the decrease of accuracy for signals with the second pure tone removed was 38.7% between Small-inter (M = 76.8%, SD = 42.3) and Large-inter condition (M = 47.1%, SD = 50) (Fig. 5C).

Two subsequent Wald tests were performed on the GLMM (*Removed pure tone*, and *Reference*) for both the Small-inter and Large-inter conditions. In Small-inter condition (Fig. 5D), the Wald test confirmed the significant impact of the *Removed pure tone* on *Accuracy* (χ2(2) = 95.3, *P* < 0.0001) and showed a significant main effect of *Reference* on *Accuracy* (χ2(3) = 8.3, *P* = 0.04) as well as a significant interaction effect between *Removed pure tone* and *Reference* (χ2(6) = 33.4, *P* < 0.0001). A post-hoc Holm’s test on this interaction revealed that significant differences between the removed pure tones were not consistent across references. Indeed, signals with the lowest and the second pure tones removed were significantly better discriminated than signals with the highest pure removed for 210+270+330 Hz and 150+210+330 Hz references (n = 1152, *P*-values < 0.0001) (Fig. 5D). Nonetheless, the removal of the lowest pure tone for the 150+210+270 Hz references and the second pure tone for the 150+270+330 Hz references were not significantly different from the removal of the highest pure tone (n = 1152, *P*-values < 0.001). In the Large-inter condition, the Wald test showed the significant impact of the *Removed pure tone* on *Accuracy* (χ2(2) = 149.4, *P* < 0.0001) as well as a significant interaction effect between *Removed pure tone* and *Reference* (χ2(6) = 23.8, *P* = 0.0005). However, no effect of the *Reference* on *Accuracy* was observed (χ2(3) = 6.7, *P* = 0.082). A post-hoc Holm’s test on *Removed pure tone* revealed that target signals with the lowest pure tone removed were significantly easier to discriminate than signals with the second pure tone removed (*P* < 0.0001), which were significantly easier to discriminate than those with the highest pure tone removed (*P* = 0.02) as shown on Figure 5C.

#### Four-tones

Only a Wald test with *Removed pure tone* and *Inter-frequency interval* as *independent variables* was performed since four-tones signals feature only one reference. The test showed a significant main effect of *Removed pure tone* on *Accuracy* (χ2(3) = 60.2, *P* < 0.0001). A post-hoc Holm’s test on *Removed pure tone* revealed that target signals with the lowest pure tone (M = 73.4%, SD = 44.3) and second pure tone (M = 65.6%, SD = 47.6) removed were significantly better discriminated than target signals with the third pure tone removed (M = 51.6%, SD = 50.1) (n = 768, *P*-values < 0.01). In turn, the third pure tone removed is significantly better discriminated than target signals with the highest pure tone removed (M = 33.9%, SD = 47.4) (n = 768, *P* < 0.001). The Wald test also showed a significant main effect of *Inter-frequency interval* on *Accuracy* (χ2(1) = 6.4, *P* = 0.011). A post-hoc Holm’s test on *Inter-frequency interval* revealed that participants were significantly more accurate in the Small-inter condition (M = 60.4%, SD = 49) than in the Large-inter condition (M = 51.8%, SD = 50) (n = 768, *P* = 0.009). The Wald test also showed a significant interaction effect between *Removed pure tone* and *Inter-frequency interval* (χ2(3) = 21.5, *P* < 0.0001). In the Small-inter condition (Fig. 5E), only the signal with the highest pure tone removed was significantly less accurately discriminated than the other signals (Holm’s test: n = 768, *P*-values < 0.05). In the Large-Inter condition (Fig. 5F), the differences between the signals were all significant with a steady decrease of accuracy as the position of the removed pure tone increased (Holm’s test: n = 768, *P*-values < 0.01).

Overall, the results from Experiment 2 are in line with those from Experiment 1; participants were generally more accurate at discriminating target signals when the lowest pure tone was removed compared to the other pure tones. The results also suggest an effect of the frequency interval between the tones since participants were surprisingly more accurate discriminating signals with small frequency intervals. All statistical effects of Experiment 2 are depicted in Table 3.

**TABLE 3.**
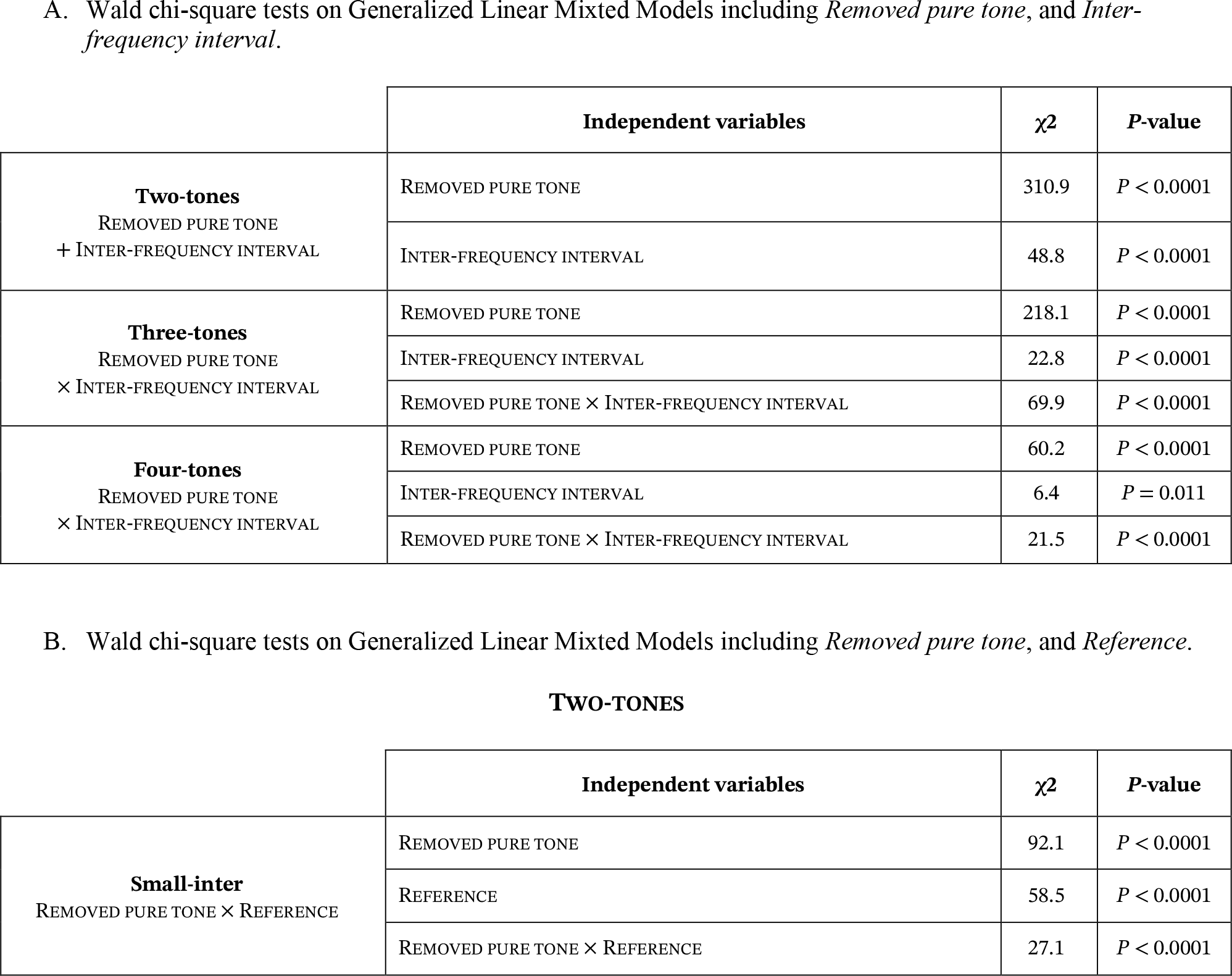
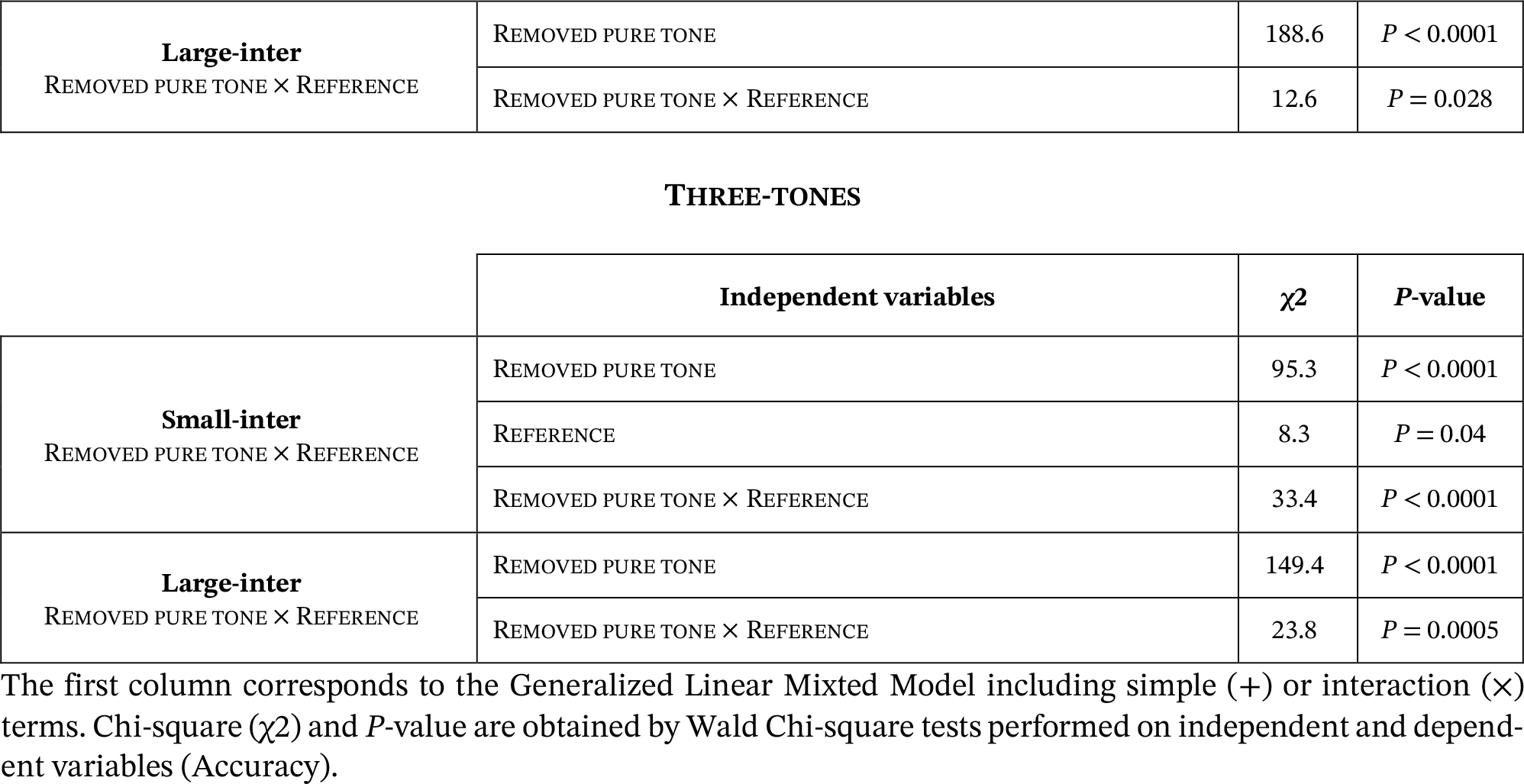
Wald chi-square tests in Experiment 2.

| | Independent variables | $\chi^2$ | P-value |
| --- | --- | --- | --- |
| **Two-tones**<br>REMOVED PURE TONE<br>+ INTER-FREQUENCY INTERVAL | REMOVED PURE TONE | 310.9 | $P < 0.0001$ |
| | INTER-FREQUENCY INTERVAL | 48.8 | $P < 0.0001$ |
| **Three-tones**<br>REMOVED PURE TONE<br>× INTER-FREQUENCY INTERVAL | REMOVED PURE TONE | 218.1 | $P < 0.0001$ |
| | INTER-FREQUENCY INTERVAL | 22.8 | $P < 0.0001$ |
| | REMOVED PURE TONE × INTER-FREQUENCY INTERVAL | 69.9 | $P < 0.0001$ |
| **Four-tones**<br>REMOVED PURE TONE<br>× INTER-FREQUENCY INTERVAL | REMOVED PURE TONE | 60.2 | $P < 0.0001$ |
| | INTER-FREQUENCY INTERVAL | 6.4 | $P = 0.011$ |
| | REMOVED PURE TONE × INTER-FREQUENCY INTERVAL | 21.5 | $P < 0.0001$ |

TWO-TONES
| | Independent variables | $\chi^2$ | P-value |
| --- | --- | --- | --- |
| **Small-inter**<br>REMOVED PURE TONE × REFERENCE | REMOVED PURE TONE | 92.1 | $P < 0.0001$ |
| | REFERENCE | 58.5 | $P < 0.0001$ |
| | REMOVED PURE TONE × REFERENCE | 27.1 | $P < 0.0001$ |
| **Large-inter**<br>REMOVED PURE TONE × REFERENCE | REMOVED PURE TONE | 188.6 | $P < 0.0001$ |
| | REMOVED PURE TONE × REFERENCE | 12.6 | $P = 0.028$ |

THREE-TONES
| | Independent variables | $\chi^2$ | P-value |
| --- | --- | --- | --- |
| **Small-inter**<br>REMOVED PURE TONE × REFERENCE | REMOVED PURE TONE | 95.3 | $P < 0.0001$ |
| | REFERENCE | 8.3 | $P = 0.04$ |
| | REMOVED PURE TONE × REFERENCE | 33.4 | $P < 0.0001$ |
| **Large-inter**<br>REMOVED PURE TONE × REFERENCE | REMOVED PURE TONE | 149.4 | $P < 0.0001$ |
| | REMOVED PURE TONE × REFERENCE | 23.8 | $P = 0.0005$ |

The first column corresponds to the Generalized Linear Mixed Model including simple (+) or interaction (×) terms. Chi-square ( $\chi^2$ ) and P-value are obtained by Wald Chi-square tests performed on independent and dependent variables (Accuracy).

### Position of the removed pure tone

Results from the two experiments suggest a significant influence of the removed pure tone’s position within the frequency spectrum of the reference. To probe more thoroughly the role of this parameter, additional analyses taking into account the frequency of the removed pure tone were performed. Since similar response patterns were observed in experiments 1 and 2 despite different frequency ranges, we tested the ratio between the removed pure tone and the lowest pure tone of the original signal (F/F_0_). This metric is also consistent with Weber’s law in which the difference limen is typically expressed as a ratio. To that end, all occurring ratios within the two experiments were considered and the individual data points with identical ratio were averaged (Fig. 6). In addition, the three levels of complexity of our study were computed separately. These data were then fitted with four plausible functions: linear, exponential, logarithmic, and sigmoidal (Fig. 6). The best goodness of fit between participants’ accuracy and the F_0_/F ratio was achieved by a sigmoidal curve for each level of complexity (Two-tones, R^2^ = 0.81; Three-tones, R^2^ = 0.74, Four-tones, R^2^ = 0.80), which matches the psychophysical expectation (Table 4). Moreover, a sum-of-squares statistical test showed that the curves fitting the three complexity levels were not significantly different (F (6,102) = 0.47, *P* = 0.82). Thus, a sigmoidal psychometric function was fitted to the entire dataset (R^2^ = 0.78) and the 66.7% just noticeable difference corresponded to a ratio of 1.47. Taken together, these results suggest that in the frequency range of our study, the discriminability of a missing tone is well predicted by its ratio with the lowest pure tone of the original signal. Interestingly, no significant effect of complexity was observed suggesting that the number of frequencies in the original signal does not affect the saliency of the removal.

**TABLE 4.**
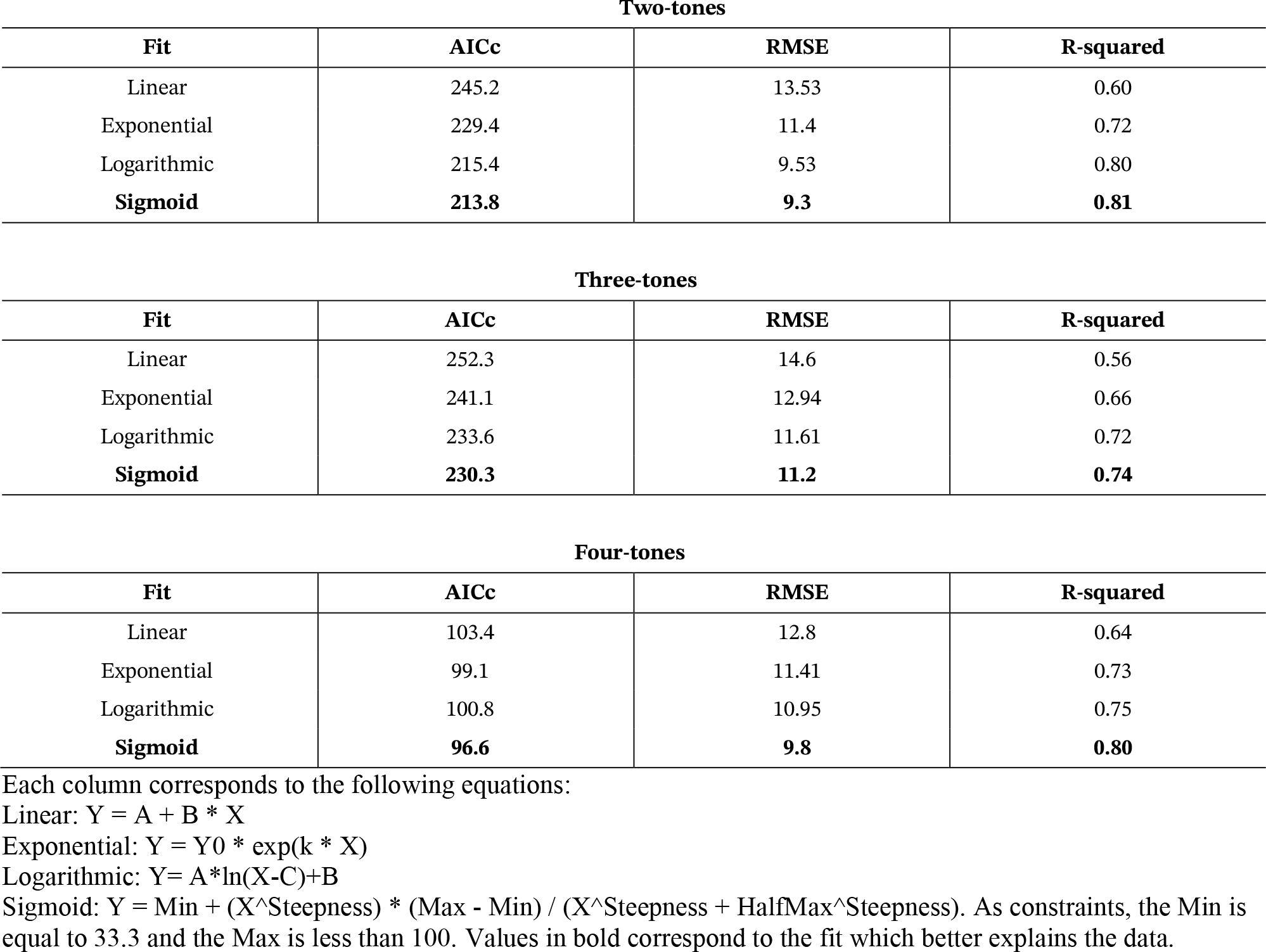
fit selection based on akaike information criterion cumulative (AICc) on the accuracy as a function of ratio.

| Two-tones |  |  |  |
| --- | --- | --- | --- |
| Fit | AICc | RMSE | R-squared |
| Linear | 245.2 | 13.53 | 0.60 |
| Exponential | 229.4 | 11.4 | 0.72 |
| Logarithmic | 215.4 | 9.53 | 0.80 |
| **Sigmoid** | **213.8** | **9.3** | **0.81** |

| Three-tones |  |  |  |
| --- | --- | --- | --- |
| Fit | AICc | RMSE | R-squared |
| Linear | 252.3 | 14.6 | 0.56 |
| Exponential | 241.1 | 12.94 | 0.66 |
| Logarithmic | 233.6 | 11.61 | 0.72 |
| **Sigmoid** | **230.3** | **11.2** | **0.74** |

| Four-tones |  |  |  |
| --- | --- | --- | --- |
| Fit | AICc | RMSE | R-squared |
| Linear | 103.4 | 12.8 | 0.64 |
| Exponential | 99.1 | 11.41 | 0.73 |
| Logarithmic | 100.8 | 10.95 | 0.75 |
| **Sigmoid** | **96.6** | **9.8** | **0.80** |

Each column corresponds to the following equations:
Linear: $Y = A + B * X$
Exponential: $Y = Y_0 * \exp(k * X)$
Logarithmic: $Y = A * \ln(X - C) + B$
Sigmoid: $Y = \text{Min} + (X^{\text{Steepness}} * (\text{Max} - \text{Min}) / (X^{\text{Steepness}} + \text{HalfMax}^{\text{Steepness}}))$ . As constraints, the Min is equal to 33.3 and the Max is less than 100. Values in bold correspond to the fit which better explains the data.

**Fig. 6.**
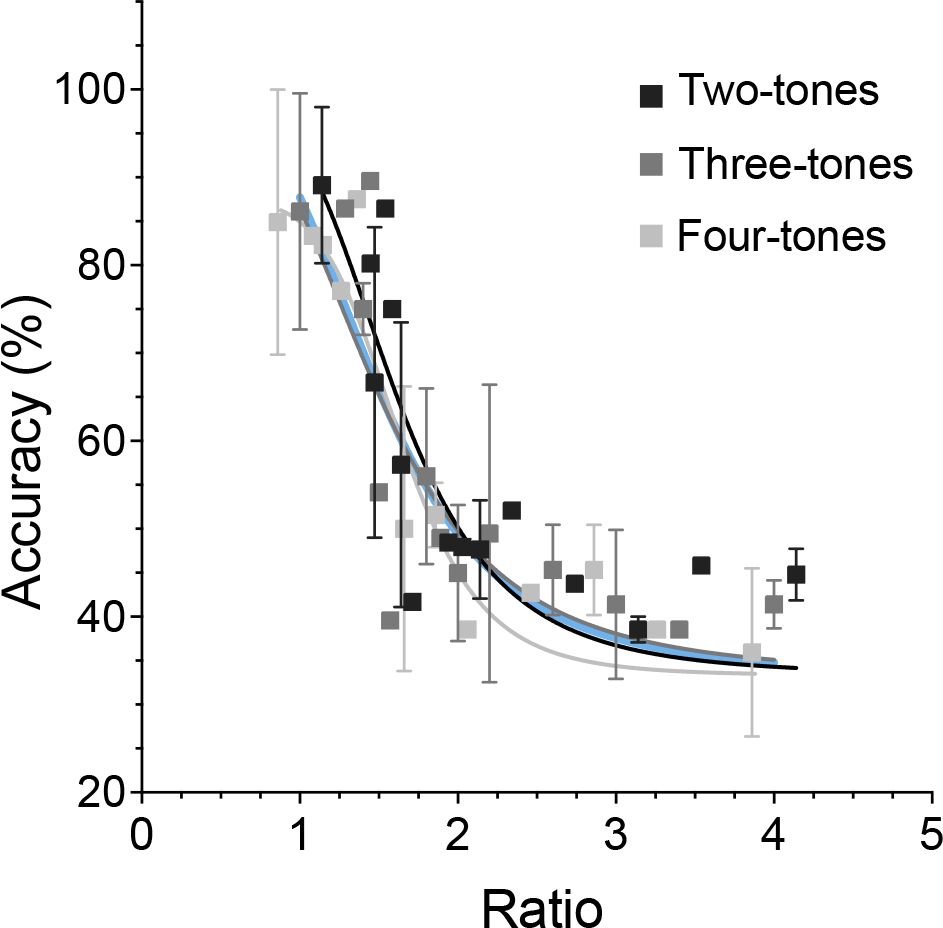
Accuracy (%) (Mean and standard error) for all the comparisons with a given ratio between the removed pure tone and the lowest pure tone of the complex signal. Each data point represents a ratio value encountered in the study. Lines correspond to a sigmoid fit for each possible complexity level of the reference signals. The blue line corresponds to the global fit for all data sets.

## Discussion

Our study investigated the potential effect of harmonicity, frequency composition, and complexity on whether removing one frequency from a complex signal would be noticeable. The obtained results showed a great variability in performance as a function of these factors. Predominantly, the results showed a significant impact of the removed pure tone’s position on discrimination. In most conditions, the lower the frequency of the removed pure tone, the more noticeable its absence. However, when a comparatively smaller relative distance between the signal’s frequencies was tested in the second experiment, the prominence of the lowest frequency decreased compared to the other constitutive frequencies of the signal. Taken together, the results of the two experiments suggest that noticing the removal of a pure tone from a complex signal does not rely on the pure tone’s position only but also on whether its frequency is close to the lower end of the signal’s spectrum. Still, it is likely that other sensory mechanisms exist since for the 120+180+240/135+195+255 Hz signals in Experiment 1, the removal of the second pure tone was noticed with significantly higher accuracy. Interestingly, no specific effect of harmonicity was observed when participants completed the task with harmonic reference signals compared to non-harmonic ones. In addition, the psychometric analysis did not show any significant effect of the complexity of the reference signal and this, regardless of the position of this frequency in the signal’s spectrum and the experiment. The dominant role of the fundamental has been pointed out in touch (20) and audition (28) but our study suggests that, more generally, the lower end of the spectrum is favored by touch following a psychometric function.

### Comparison with previous findings

The lack of influence of the harmonicity of the signal is somewhat surprising, given that the harmonicity and consonance of vibrotactile signals have been shown to impact perception (22, 23, 29). However, previous studies compared signals of identical complexity while the signals in our experiments differ by a missing frequency, which probably introduces stronger sensory phenomena than harmonicity. Additionally, differences in diharmonic signals have been shown to be slightly more difficult to distinguish than in pure tones (16, 17), while the results of our study did not show an impact of complexity. Based on the frequency selectivity of psychophysical channels (25, 30, 31) and their independence when it comes to masking (32–34), one could have expected the effect of removing pure tones to differ depending on whether or not they belong to the same tactile channel. However, the experimental results, which encompass a large frequency range from 60 Hz to 600 Hz, suggest that the main tactile channel activated by the removed pure tone has little impact on how its removal is perceived. The observed human sensitivity to removed frequencies is also intriguing since it does not follow the classical relationship of tactile sensitivity in which the detection threshold is lowest for high frequencies around 300 Hz (30, 35). However, our stimuli were chosen to be largely over the human threshold for detecting vibrations, which might explain why tactile sensitivity does not play a role.

Additional analyses that focus deeper on the frequency characteristics of the removed pure tone showed that differences in the ratio between the removed pure tone and the lowest pure tone accurately predicts the capacity to detect a missing pure tone. Moreover, these analyses showed that a complex signal with N-1 frequencies can be indistinguishable from a simpler signal featuring N frequencies depending on where the removed frequency lies within the signal’s spectrum. These results extend existing findings that the lower frequencies of two-tones complex signals are more perceptually salient (20).

### Physical and cognitive mechanisms underlying vibrotactile perception

The dependence on frequency rather than intensity of our perceptual results is in line with the spectral model of vibrotactile perception, which suggests that the percept of a complex signal comes from the sum of the relative activations of the frequency-tuned minichannels in the Pacinian sensitivity range. However, this model stems from the Pacinian activity while the supra-threshold nature of the stimuli in this study suggests that the delivered vibrations were encoded by most of the tactile receptors, and it is also known that several fundamental tactile cues are processed in the somatosensory cortex (36). Similarly, Russo et al. (23) have supported the idea that frequency-tuned populations of mechanoreceptors filter complex tones into their component frequencies. Moreover, frequency-mediated sensory mechanisms are also supported by evidence that vibrotactile frequency perception relies on the duration of individual inter-tap intervals rather than the average firing rate (37, 38), and this even for complex frequency compositions or signals without fixed periodicity. Although the perceptual cues used by participants remain to be elucidated, this could explain the dominant role of the lower frequencies, which are responsible for the larger and possibly most salient intervals between vibrotactile bursts. It is also possible that this pattern also results from masking phenomena. Indeed, previous research suggests that narrow-band noise in the Pacinian channel has a slightly weaker masking effect on the detection of a 20 Hz sinusoidal signal than Non-Pacinian noise on a 200 Hz signal (32). These results do not fully follow the predictions based on the separation of tactile channels and might also relate to the frequency selectivity mechanisms observed in our study. This suggests a potential interplay of physical, sensory, and cognitive mechanisms.

### Perspectives and Significance

Our study shows a large unbalance in the perceptual weight of frequencies within complex signals up to four tones. The dominance of the fundamental pure tone of a diharmonic signals has already been suggested (20) but our results additionally show that it is the value of the frequency within the complex signal’s spectrum that matters even if it is the second or third largest frequency. The number of frequencies within the signal does not seem to influence the perceptual value of a given pure tone. To investigate further the generality of these findings, it would be interesting to extend the experiments to frequencies under 60 Hz, more variable inter-frequency intervals, and stimuli that are closer to the human sensory threshold. Although non-Pacinian channels were also targeted in Experiment 1 with the 60 and 75 Hz, these frequencies also activated the Pacinian channel. Testing frequencies around 10-30 Hz that dissociate the activation of the psychophysical tactile channels (25, 30, 31) would enable to thoroughly investigate their potential role in the perception of complex vibrotactile waves. Extending the comparisons to complex signals with a larger number of frequencies and a different organization of the frequency spectrum would also be insightful. Measuring the detection threshold of complex signals could also be useful for predicting discrimination (19) as well as determining the masking effects elicited by the interplay between frequencies (32–34). Finally, we chose to equalize the amplitude of the constitutive pure tones in peak acceleration, but it would also be interesting to evaluate the same task with pure tones of various amplitudes or equalized in perceived intensity. The present study paves the way for a better understanding of the main factors affecting perception of complex vibrotactile signals and the development of a more general psychophysical model. It also opens new perspectives regarding the design of vibrotactile signals in industry applications since it shows that complex vibrotactile signals can be largely simplified without any perceptual change for the user.

**Current adresses :** Institut des Systèmes Intelligents et de Robotique, Sorbonne Université, Paris, 75005.

## Data availability

The data associated with this study are available to the scientific community to ensure transparency, reproducibility, and further exploration of the findings presented in this article.

Researchers interested in accessing the data can find it in the following locations (in viewable condition only): https://www.dropbox.com/scl/fo/90k0sil2jxv8sriqg7at1/h?rlkey=58tapjxu59aczb0ib6netbsyh&dl=0

## Primary Data Repository

All primary experimental data are archived and publicly available on *Data* folder.

For any inquiries or requests related to the data, please feel free to contact the corresponding author at.

## Acknowlegments

We are grateful for the kind participation of all participants. We thank Dr. Georges Daher for his technical support.

## Authors contributions

T-L S.L, G.B, E.V, M.A, and D.G conceived and designed research, T-L S.L performed experiments, T-L S.L, G.B, M.A. and D.G analyzed data and interpreted the results of experiments, T-L S.L, G.B., and D.G. prepared figures, and drafted manuscript, T-L S.L, G.B, M.A, and D.G edited and revised manuscript, T-L S.L, G.B, E.V, M.A, and D.G approved final version of manuscript.

